# Quantification of NAD^+^ T_1_ and T_2_ relaxation times using downfield ^1^H MRS at 7 T in human brain in vivo

**DOI:** 10.1101/2024.02.27.582276

**Authors:** Sophia Swago, Neil E. Wilson, Mark A. Elliott, Ravi Prakash Reddy Nanga, Ravinder Reddy, Walter R. Witschey

## Abstract

**Introduction:** The purpose of this study was to use a single-slice spectrally-selective sequence to measure T_1_ and T_2_ relaxation times of NAD^+^ proton resonances in the downfield ^1^H MRS spectrum in human brain at 7 T in vivo and assess the propagation of relaxation time uncertainty in NAD^+^ quantification.

**Methods:** Downfield spectra from 7 healthy volunteers were acquired at multiple echo times in all subjects to measure T_2_ relaxation, and saturation recovery data were to measure T_1_ relaxation. The downfield acquisition used a spectrally-selective 90° sinc pulse for excitation centered at 9.1 ppm with a bandwidth of 2 ppm, followed by a 180° spatially-selective Shinnar-Le Roux refocusing pulse for localization. For the multiple echo experiment, spectra were collected with echo times ranging from 13 to 33 ms. For the saturation recovery experiment, saturation was performed prior to excitation using the same spectrally-selective sinc pulse as was used for excitation. Saturation delay times (TS) ranged from 100 to 600 ms. Uncertainty propagation analysis was performed analytically and with Monte Carlo simulation.

**Results:** The mean ± standard deviation of T_1_ relaxation times of the H2, H6, and H4 protons were 152.7 ± 16.6, 163.6 ± 22.3, and 169.9 ± 11.2 ms, respectively. The mean ± standard deviation of T_2_ relaxation times of the H2, H6, and H4 protons were 32.5 ± 7.0, 27.4 ± 5.2, and 38.1 ± 11.7 ms, respectively. The mean R^2^ of the H2 and H6 T_1_ fits were 0.98. The mean R^2^ of the H4 proton T_1_ fit was 0.96. The mean R^2^ of the T_2_ fits of the H2 and H4 proton resonances were 0.98, while the mean R^2^ of the T_2_ fits of the H4 proton was 0.93. The relative uncertainty in NAD^+^ concentration due to relaxation time uncertainty was 8.5%-11%.

**Conclusion:** Using downfield spectrally-selective spectroscopy with single-slice localization, we found NAD^+^ T_1_ and T_2_ relaxation times to be approximately 162 ms and 32 ms respectively in the human brain in vivo at 7 T.

## Introduction

Nicotinamide adenine dinucleotide (NAD^+^) is a key metabolite involved in many cellular processes. In energy metabolism, NAD^+^ is reduced to NADH in the tricarboxylic acid cycle, and NADH is in turn oxidized to NAD^+^ in the electron transport chain during oxidative phosphorylation. It additionally serves as a cofactor for enzymes that carry out post-translational modification, including poly-ADP-ribose polymerases (PARPs) and sirtuins, which are involved in cell signaling and survival (1). In recent years, NAD^+^ has gained much interest as a biomarker and therapeutic target for its role in aging and disease; NAD^+^ decline has been observed in the aging brain, as well as in neurodegenerative and cardiovascular diseases (2-5). NAD^+^ has emerged as an attractive therapeutic target in part due to the availability of NAD^+^ precursors, such as nicotinamide riboside (NR) and nicotinamide mononucleotide (NMN), which may be able to modulate NAD^+^ levels in vivo (6-8). Current methods used to measure NAD^+^ include invasive techniques such as high-performance liquid chromatography (HPLC) using tissue samples, and noninvasive methods such as 31P magnetic resonance spectroscopy and fluorescent imaging (9-11). However, HPCL is limited by the availability of tissue that can be biopsied as well as metabolite stability, while fluorescent imaging is not able to detect NAD^+^ in deep organs. ^31^P MRS suffers from decreased sensitivity due to its lower gyromagnetic ratio in comparison to ^1^H MRS and requires specialized MR coils tuned to the phosphorous resonances.

Recently, there have been advances made in the detection of NAD^+^ resonances in the downfield (>4.7 ppm) region of the ^1^H MR spectrum (12-16). However, there are several challenges that hinder the measurement of downfield metabolites using conventional MRS experiments: the metabolites with downfield resonances have inherently low concentrations (0.1-10 mM); perturbation of the water protons leads to downfield metabolite signal loss through either chemical exchange or cross-relaxation with water (13,17,18); there may be significant overlap of resonances; and many additionally have short T_2_ relaxation times (20-40 ms at 7 T) (19) requiring the use of short echo time sequences. Downfield spectroscopy techniques therefore avoid the use of water suppression through either spectrally-selective excitation or metabolite cycling (20), which alternately inverts the upfield and downfield regions. Downfield spectroscopy techniques also benefit from ultrahigh field strengths (i.e. 7 T), which provides increased sensitivity and chemical dispersion leading to better discrimination of resonance peaks.

An important aspect of accurate quantification of metabolite concentrations using MRS is correction for T_1_ and T_2_ relaxation effects on the resultant spectra. While MacMillan et al. and Fichtner et al. measured T_1_ and T_2_ relaxation times of downfield metabolites at 3 T and 7 T in human brain, NAD^+^ was undetectable in these studies (19,21). Borbath et al. found the T_2_ relaxation time of NAD^+^ to be 30 ms at 9.4 T in human brain but calculated this value from spectra summed across all subjects (22). Finally, de Graaf et al. determined that the T_1_ and T_2_ of NAD^+^ were 280 ms and 60 ms, respectively, in rat brain in situ at 11.7 T (13). To date, all studies that have reported NAD^+^ concentration in human brain have used extrapolated T_1_ and T_2_ times derived from de Graaf’s measurements, but this extrapolation is a potential source of error in NAD^+^ concentration measurements (12,14,15).

In this work, we report the T_1_ and T_2_ relaxation times of downfield NAD^+^ resonances measured using downfield spectrally-selective spectroscopy with multiple saturation delays and echo times in the human brain in vivo at 7 T and measure absolute NAD^+^ concentration in a subset of participants. Additionally, we investigate the propagation of uncertainty from relaxation time measurements to metabolite concentration quantification using both analytical and Monte Carlo methods.

## Methods

### Participants

All participants were scanned in accordance with the local Institutional Review Board protocols after giving written, informed consent. T_1_ saturation recovery and T_2_ decay data acquired using ^1^H downfield MRS were collected on 7 healthy human volunteers without a history of neurological or psychiatric disease (3 male and 4 female) between the ages of 25 and 30.

### In vivo magnetic resonance spectroscopy

Subjects were scanned at 7 T (Terra, Siemens Healthcare, Erlangen, Germany) using a 32-channel RF head coil (Nova Medical, Wilmington, MA, USA). The spectroscopy slice was positioned using a localizer. The slice was oriented in the axial plane through the brain and subsequently obliqued to avoid the frontal sinus and had a thickness of 40 mm. A typical slice localization is shown in Figure 1A. Manual shimming over the slice was performed, with an average reference water peak full-width half-maximum (FWHM) of 22.8 ± 2.8 Hz. The downfield MRS acquisition used a spectrally-selective 90° sinc pulse for excitation centered at 9.1 ppm with a bandwidth of 2 ppm, followed by a 180° spatially-selective Shinnar-Le Roux refocusing pulse for slice localization (Fig. 1B). The NAD^+^ resonances of interest arise from the H2, H6, and H4 protons of nicotinamide moiety at 9.3, 9.1, and 8.9 ppm, respectively (Fig. 1C) (13).

**Figure 1.**
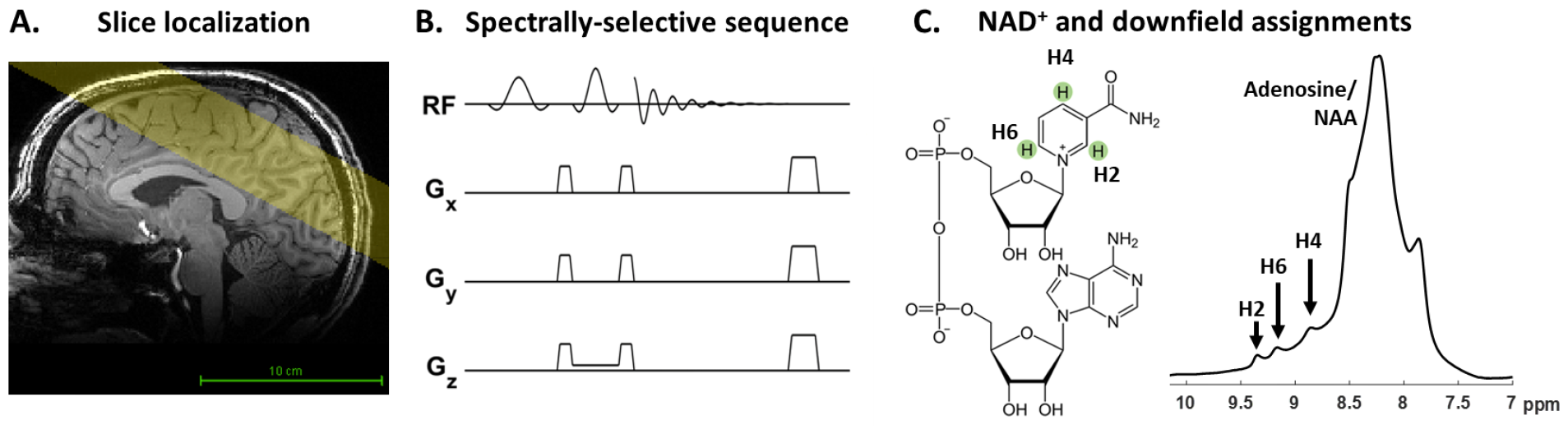
A) Representative example of slice localization overlaid on a T_1_-weighted image. B) Schematic of the single-slice spectrally-selective sequence. C) The chemical structure of NAD^+^ and corresponding downfield ^1^H MRS peak assignments.

In the T_1_ saturation recovery experiment, saturation was performed prior to excitation using the same spectrally-selective sinc pulse as was used for excitation. Spectra were acquired with a range of saturation delay times (TS) between 100and 600 ms and TR/TE = 2500/13 ms. An additional dataset without saturation but with the same TR/TE was acquired to measure the equilibrium or fully recovered magnetization (M_0_). The spectra with saturation delay times of 100 and 200 ms were acquired with 256 averages (10.7 minutes) to boost SNR; all other spectra were acquired with 128 averages (5.3 minutes). The total acquisition time of the saturation recovery experiment was approximately 37 minutes.

For the T_2_ decay experiment, spectra were collected at a range of echo times between 13 and 33 ms, with TR=1000 ms and averages = 128 (2.1 minutes). The total acquisition time of the multiple echo experiment was approximately 12 minutes. In all scan sessions, a water reference spectrum was also acquired using the same sequence used to acquire the metabolite spectra with the excitation center changed to 4.7 ppm, TR/TE = 1000/13 ms, averages = 16. In a subset of subjects (N=4), a high resolution T_1_-weighted MP2RAGE anatomical brain scan was acquired for tissue segmentation: TR: 2260 ms, TE: 3.71 ms, TI: 1100 ms, resolution: 0.9375 × 0.9375 × 1 mm.

### MRS data processing

All MRS processing was done using custom software (Matlab, The Mathworks, Natick, MA, USA). Coil channel combination was done offline for the spectroscopy datasets using the water reference raw FID data to determine channel weightings. Additionally, a channel-wise frequency correction was applied to account for B0 differences within the slice and then applied to the downfield metabolite FID data.

### MRS data fitting

All spectra were 5 Hz line broadened. The scan with the highest signal in each experiment (M_0_ in the T_1_ experiment or shortest TE in the T_2_ experiment) was fit in the time domain using Hankel singular value decomposition (HSVD) (23,24). Components found within empirically chosen constraints for peak center frequency and linewidth were assigned to the three NAD^+^ peaks at 9.3, 9.1, and 8.9 ppm. All other components were assigned to the baseline. The resonance centers and linewidths of the components that were assigned to the NAD^+^ resonances were fixed to form a spectral basis set, and a linear regression was performed to fit the amplitude and phase of the remaining datasets in the saturation recovery and multiple echo time experiments. Additional frequency correction was performed between subsequent scans within the multiple echo experiments and the saturation recovery experiment.

For fitting T_1_ relaxation time, the dataset from the saturation recovery experiment without saturation was assigned a saturation delay time of 5 s, a sufficiently long time to represent the equilibrium magnetization. For each of the three NAD^+^ peaks, the amplitudes at each saturation delay time were fit using a two-parameter model for T_1_ relaxation time: 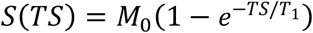.

For fitting T_2_ relaxation times, the fitted NAD^+^ amplitudes from the multiple TE experiment were first corrected for the effects of J-modulation by integrating simulated signal at each TE using the scalar couplings determined by de Graaf et al (13). The corrected amplitudes were then fit using a two-parameter model for T_2_ relaxation time: 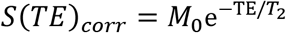.

In a subset of participants (N=4), the concentration of NAD^+^ was calculated:

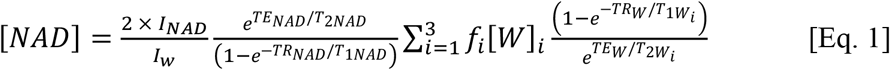

where [NAD] is the concentration on NAD^+^, I_NAD_ is the amplitude of the NAD^+^ peak, TE_NAD_ and TR_NAD_ are the echo and repetition time of the NAD^+^ downfield scan. I_W_ is the amplitude of the water peak from the water reference scan, a factor of 2 is included to account for the two protons in water, f_i_ is the fraction of brain tissue *i* (cerebral spinal fluid, gray matter, or white matter) determined from the T_1_w anatomic image, [W]_i_ is the water content of brain tissue *i*, TE_W_ and TR_W_ are the echo and repetition time of the water reference scan, and T_1Wi_ and T_2Wi_ are tissue specific water relaxation times (25). The brain segmentation to determine tissue fractions consisted of the following: N4 bias field correction (26), brain extraction (27), and FSL FAST segmentation for CSF, gray matter, and white matter (28). Only the 9.3 ppm (H2 proton) NAD^+^ peak was used to measure the NAD^+^ concentration because it had the least amount of overlap with the complex of peaks found in the 8 to 8.8 ppm range.

### Uncertainty propagation analysis

To evaluate the effect of the uncertainty of the measured T_1_ and T_2_ values on the calculation of NAD^+^ concentration using the H2 proton, we determined the propagation of the uncertainty using both analytic and Monte Carlo approaches. In the analytic approach, the uncertainty in a function outcome *F* due to the uncertainty in a parameter *X* is described by 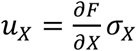 where 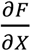 is the partial derivative with respect to parameter *X* and *σ*_*X*_ is the standard deviation of *X*. For *n* independent parameters, the combined uncertainty (estimated standard deviation) is (29):

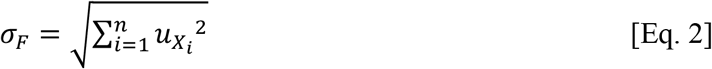

The partial derivatives of [NAD] with respect to T_1NAD_ and T_2NAD_ were derived by Instrella and Juchem (30):

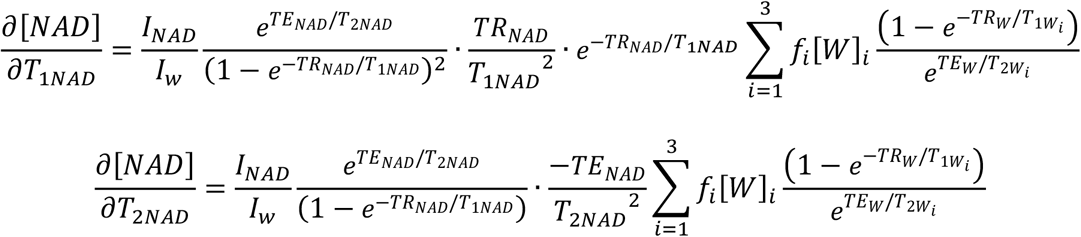

The separate influence of T_1NAD_ and T_2NAD_ relaxation times on the uncertainty was assessed by calculating the absolute value of the respective partial derivatives across a range of times (T_1_: 0-400 ms, T_2_: 0-150 ms) while all other parameters were held constant as found in Table 1. The combined uncertainty was calculated as described in Eq. 2 using the mean and standard deviations of T_1_ and T_2_ relaxation times derived from all subjects. The relative uncertainty is reported as the combined uncertainty divided by the estimate of [*NAD*] using the mean T_1_ and T_2_.

**Table 1.**
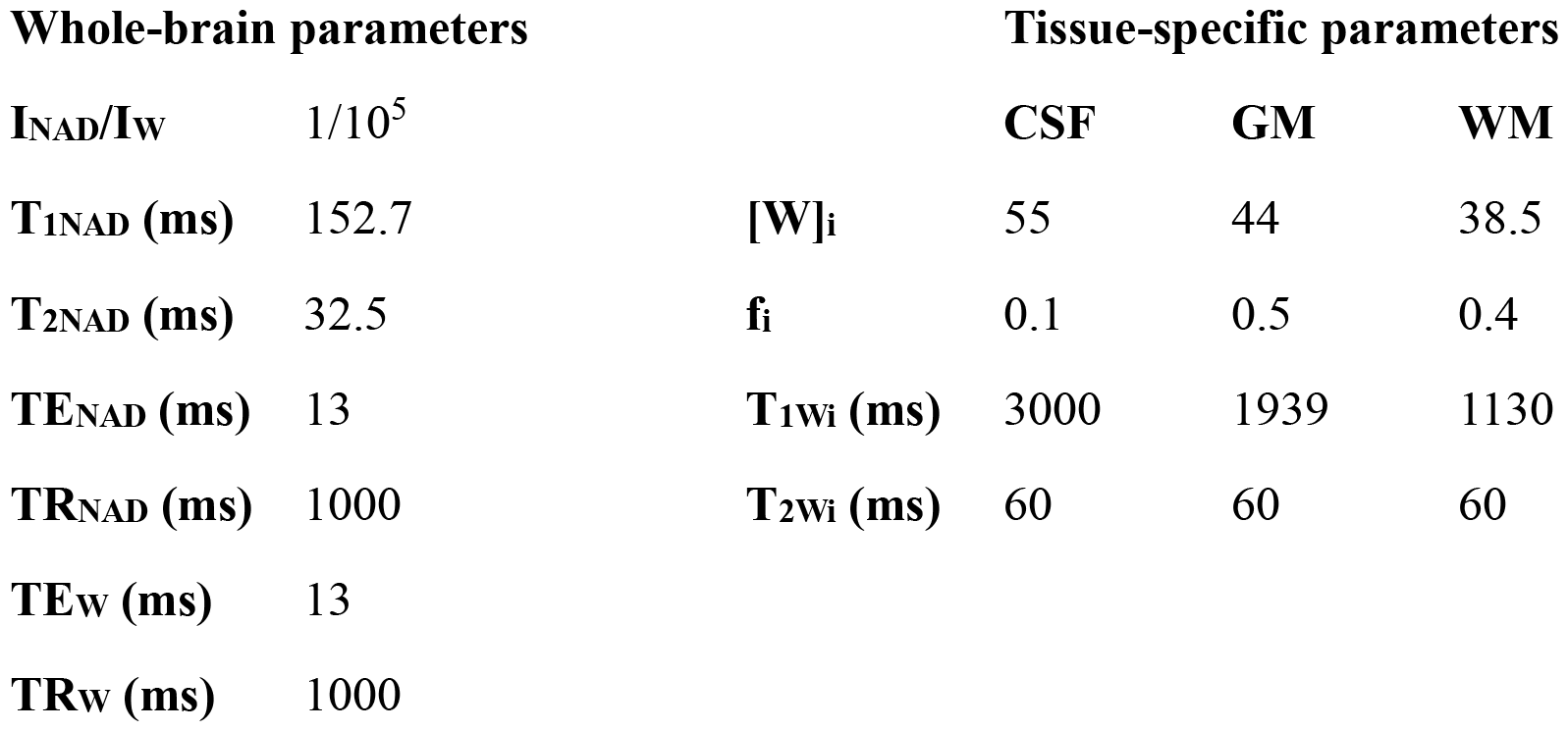
Parameter constants in NAD^+^ quantification and uncertainty analysis. T_1NAD_ and T_2NAD_ were only held constant in the individual parameter partial derivative analysis. Brain tissue water content and relaxation times from (25).

| Whole-brain parameters |  |  | Tissue-specific parameters |  |  |
| --- | --- | --- | --- | --- | --- |
| $I_{\text{NAD}}/I_{\text{W}}$ | $1/10^5$ | | CSF | GM | WM |
| $T_{1\text{NAD}}$ (ms) | 152.7 | $[\text{W}]_{\text{i}}$ | 55 | 44 | 38.5 |
| $T_{2\text{NAD}}$ (ms) | 32.5 | $f_{\text{i}}$ | 0.1 | 0.5 | 0.4 |
| $TE_{\text{NAD}}$ (ms) | 13 | $T_{1\text{Wi}}$ (ms) | 3000 | 1939 | 1130 |
| $TR_{\text{NAD}}$ (ms) | 1000 | $T_{2\text{Wi}}$ (ms) | 60 | 60 | 60 |
| $TE_{\text{W}}$ (ms) | 13 | | | | |
| $TR_{\text{W}}$ (ms) | 1000 | | | | |

For the Monte Carlo approach, 10^5^ random T_1_ and T_2_ values were drawn from normal distributions of T_1_ and T_2_ times generated using the means and standard deviations calculated across all subjects for the H2 proton. All other parameters were held constant at the same values used in the analytic analysis. These relaxation times were then used to create a distribution of metabolite concentrations. The standard deviation of this distribution is the estimated uncertainty in the concentration measurement. As in the analytic approach, the relative uncertainty is the estimated uncertainty divided by the estimated concentration calculated using the mean T_1_ and T_2_.

## Results

During HSVD fitting, a single Lorentzian was assigned to each of the peaks found at 9.1, 9.1, and 8.9 ppm, which arise from the H2, H6, and H4 protons of NAD^+^ respectively (Figure 2A). The linewidths of the NAD^+^ peaks found in the saturation recovery experiment were 25.5 ± 1.1 Hz (H2), 32.3 ± 1.2 Hz (H6), and 36.1 ± 2.4 Hz (H4); the linewidth for the fitted NAD^+^ peaks in the multiple echo experiment were 28.2 ± 5.1 Hz (H2), 34.9 ± 7.2 Hz (H6), and 39.2 ± 7.0 Hz (H4). Representative series of spectra at multiple saturation delay times and multiple echo times are shown in Figure 2B and Figure 2C.

**Figure 2.**
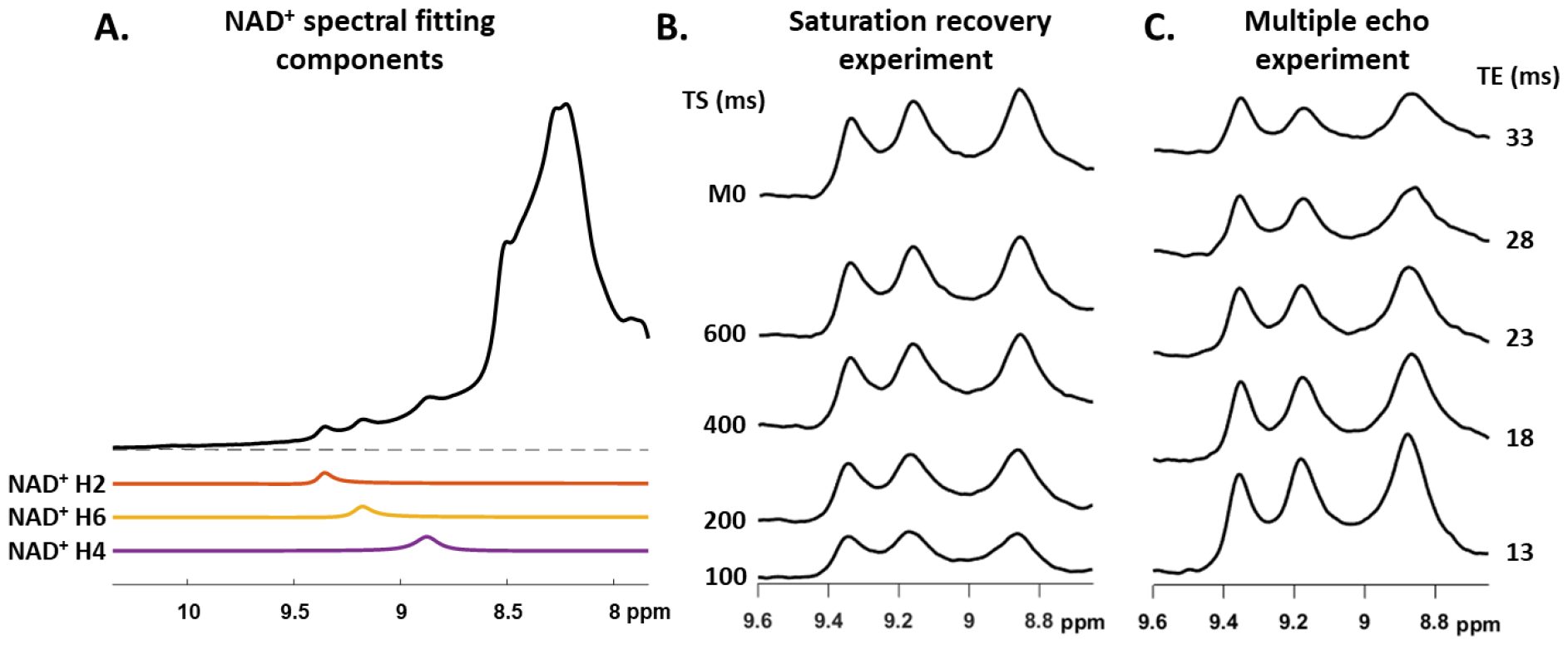
A) A measured spectrum is shown with the HSVD fit components assigned to each of the three NAD^+^ peaks. B) Representative series of spectra from a saturation recovery experiment; signal increases with increasing saturation delay time (TS). C) Representative series of spectra from a multiple echo time experiment; signal decreases with increasing echo time (TE).

Representative T_1_ and T_2_ (after J-modulation correction of amplitudes) fits from a single subject for all three NAD^+^ peaks are shown in Figures 3 and 4. The T_1_ and T_2_ relaxation time fits for all subjects are summarized in Table 2. The mean T_1_ relaxation times for each NAD^+^ peak were: 152.7 ± 12.2 ms (H2), 163.6 ± 22.3 ms (H6), and 169.9 ± 11.2 ms (H4) (Fig. 3B). The mean T_2_ relaxation times were: 32.5 ± 7.0 ms (H2), 27.4 ± 5.2 ms (H6), and 38.1 ± 11.7 ms (H4) (Fig. 4B). The quality of the relaxation time fits was assessed by coefficient of determination (R^2^). The mean R^2^ of the T_1_ fits were 0.98 ± 0.01 (H2), 0.98 ± 0.01 (H6), and 0.96 ± 0.06 (H4); for the T_2corr_ fits, mean R^2^ were: 0.98 ± 0.01 (H2), 0.98 ± 0.02 (H6), and 0.93 ± 0.07 (H4). In a subset of subjects, NAD concentration was measured from the H2 proton using Eq. 1 and was found to be 0.300 ± 0.064 mM.

**Table 2.** Fitted T_1_ and T_2_ relaxation times of all subjects and R^2^ of all fits for each of the three NAD^+^ peaks.

| $NAD^+$ proton | $T_1$ (ms) | | | $T_1 R^2$ | | |
| --- | --- | --- | --- | --- | --- | --- |
|  | H2 | H6 | H4 | H2 | H6 | H4 |
| **Subject 1** | 147.8 | 161.1 | 189.1 | 0.97 | 0.97 | 0.98 |
| **Subject 2** | 146.1 | 171.2 | 155.0 | 0.99 | 0.99 | 0.97 |
| **Subject 3** | 147.3 | 149.5 | 175.7 | 0.98 | 0.97 | 0.99 |
| **Subject 4** | 148.2 | 165.0 | 167.7 | 0.99 | 1.00 | 0.97 |
| **Subject 5** | 140.3 | 140.9 | 159.7 | 0.99 | 0.99 | 0.98 |
| **Subject 6** | 175.0 | 208.2 | 168.0 | 0.98 | 0.99 | 0.83 |
| **Subject 7** | 164.2 | 149.1 | 173.8 | 0.99 | 0.99 | 0.95 |
| **Mean+std** | 152.7±12.2 | 163.6±22.3 | 169.9±11.2 | 0.98±0.01 | 0.98±0.01 | 0.96±0.06 |

| $NAD^+$ proton | $T_2$ (ms) | | | $T_2 R^2$ | | |
| --- | --- | --- | --- | --- | --- | --- |
|  | H2 | H6 | H4 | H2 | H6 | H4 |
| **Subject 1** | 38.0 | 22.6 | 25.8 | 0.99 | 0.93 | 0.89 |
| **Subject 2** | 30.6 | 26.9 | 41.0 | 0.98 | 0.99 | 0.97 |
| **Subject 3** | 28.7 | 26.8 | 47.3 | 0.96 | 0.98 | 0.98 |
| **Subject 4** | 34.5 | 34.7 | 47.7 | 0.99 | 0.99 | 0.91 |
| **Subject 5** | 44.2 | 33.9 | 28.2 | 0.95 | 0.97 | 0.79 |
| **Subject 6** | 23.0 | 21.0 | 24.6 | 0.99 | 0.99 | 0.98 |
| **Subject 7** | 28.9 | 26.2 | 52.3 | 0.99 | 0.98 | 0.98 |
| **Mean+std** | 32.5±7.0 | 27.4±5.2 | 38.1±11.7 | 0.98±0.01 | 0.98±0.02 | 0.93±0.07 |

**Figure 3.**
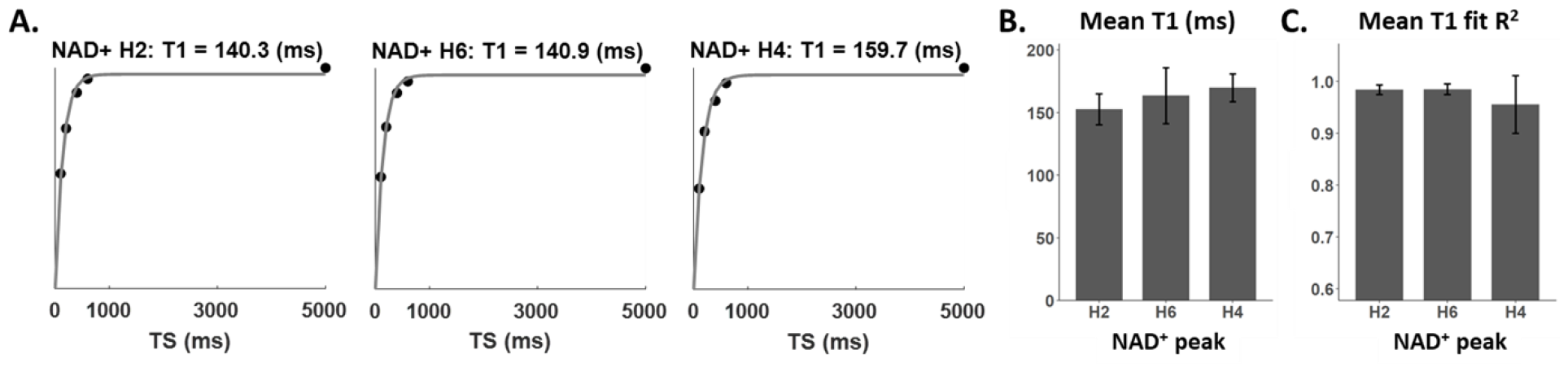
A) Representative T_1_ saturation recovery curves from a single subject showing the data (points) and the modeled fit (lines) for each of the NAD^+^ peaks. B) The mean T_1_ relaxation time and C) the mean R^2^ of the T_1_ fit across all subjects for each NAD^+^ peak.

**Figure 4.**
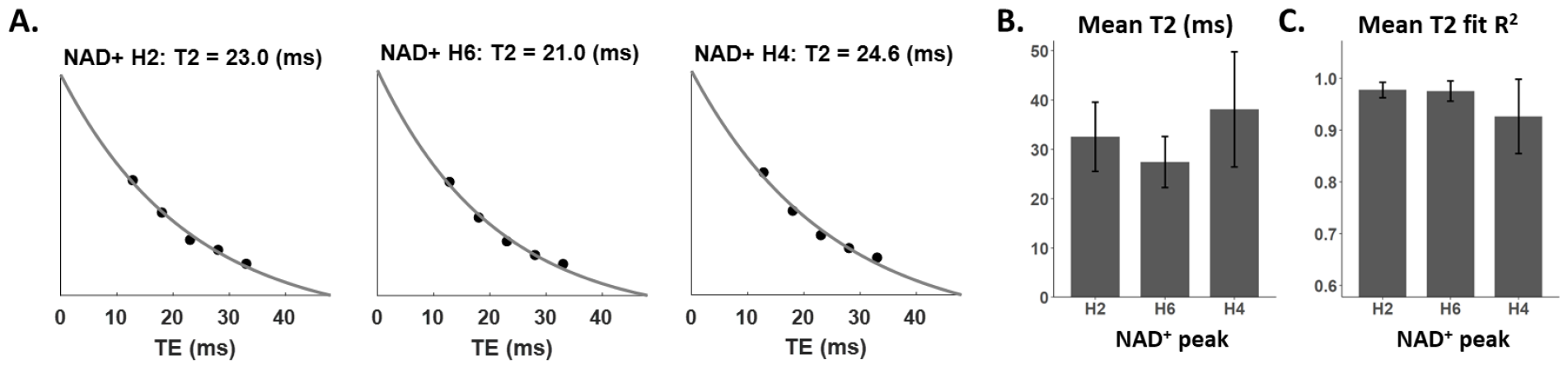
: A) Representative T_2_ exponential decay fits after amplitude correction for J-modulation effects from a single subject. B) The mean T_2_ relaxation time and C) the mean R^2^ of the T_2_ fit across all subjects for each NAD^+^ peak.

Uncertainty propagation analysis was carried out for the H2 NAD^+^ proton. In the analytic model, the variance of its T_1_ and T_2_ relaxation times was first assessed separately. The partial derivative of concentration with respect to each relaxation time parameter over a range of relaxation times is shown in Figure 5A. The magnitude of the partial derivative with respect to T_1_ relaxation time was smaller than that with respect to T_2_ relaxation time (2.8·10^−5^ mM/ms vs 5.7·10^−3^ mM/ms). The absolute combined uncertainty was 0.040 mM, and the relative uncertainty was 8.5%. The Monte Carlo derived concentration estimates are shown in Figure 5B. The resulting distribution is skewed right due to the nonexponential nature of the T_1_ and T_2_ terms in the equation to calculate concentration. The standard deviation of the distribution, representing the estimated uncertainty due to the relaxation time parameters, was 0.054 mM and the estimated fractional uncertainty was 11.5%.

**Figure 5.**
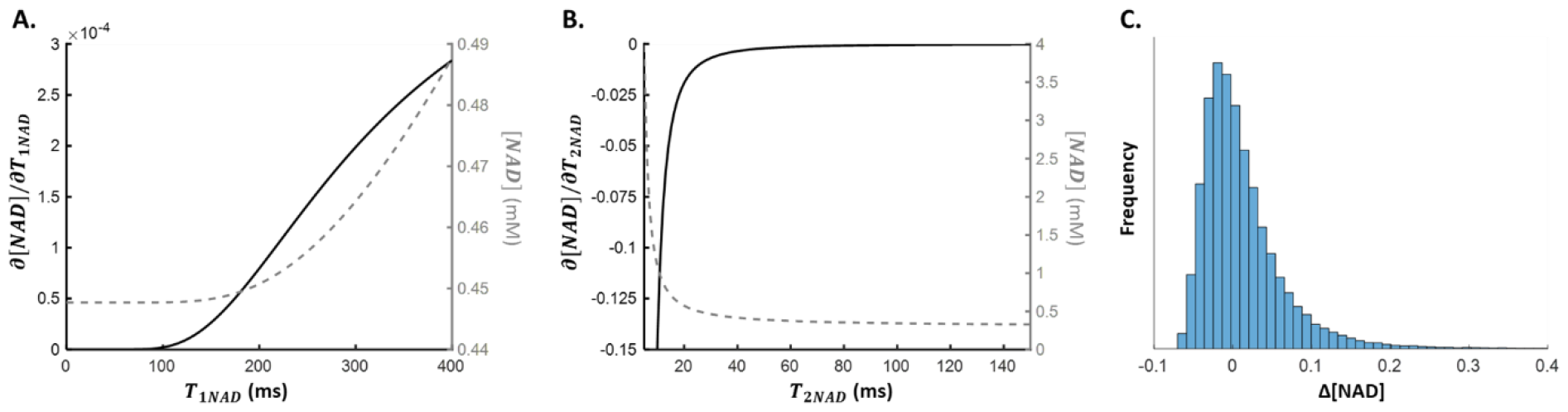
A) Analytic solution to ∂ [*NAD*]*/*∂*T*_*1NAD*_ (solid black line) and the corresponding [*NAD*] quantification (dotted gray line) as a function of T_1NAD_. B) Analytic solution to ∂ [*NAD*]*/*∂*T*_2*NAD*_ (solid black line) and the corresponding [NAD] quantification (dotted gray line) as a function of T_2NAD_. C) Monte Carlo distribution of estimated absolute concentration of NAD^+^ normalized by the true concentration.

## Discussion

The aim of this study was to determine the T_1_ and T_2_ relaxation times of downfield NAD^+^ proton resonances in the human brain in vivo at 7 T using a spectrally selective single-slice sequence. Previous studies at 3 T, 7 T, and 9.4 T have investigated relaxation times of downfield metabolites in the brain; in general, it was found that downfield metabolites exhibit shorter T_1_ and T_2_ times than upfield metabolites (19,21,31). However, the NAD^+^ resonances at 9.3, 9.1, and 8.9 ppm were not visible in these experiments. These studies used stimulated echo acquisition mode (STEAM) single voxel spectroscopy either in combination with metabolite cycling or with water suppression and did not have sufficient SNR to detect NAD^+^. In comparison, our study used a 40 mm slice, and a spin-echo sequence without water suppression to enhance NAD^+^ detection, allowing us to fit the three NAD^+^ resonances at a variety of echo times and saturation delay times.

We found the T_1_ and T_2_ of NAD^+^ to be 162 ms and 32 ms respectively. This T_1_ is shorter than that found by de Graaf in rat brain at 11.7 T (13). That study found the T_1_ of NAD^+^ to be 280 ms using a spectrally-selective inversion recovery experiment. This is in line with the expectation that T_1_ decreases at lower field strengths. However, though it is expected that T_2_ decreases with field strength, de Graaf determined the T_2_ of NAD^+^ to be 60 ms at 11.7 T while the T_2_ determined in the present study was notably shorter. Nonetheless, the T_2_ time we found was similar to the T_2_ of NAD^+^ found by Borbath et al., who measured it to be 30.3 ms at 9.4 T in human brain, though this was measured after summing the spectra of all subjects (22). We additionally found that the T_1_ and T_2_ of the H2 proton of NAD^+^ had low variance across subjects and high fit reliability as determined by the fit R^2^ in comparison to the H6 and H4 protons. This is likely due to the larger amount of overlap the H6 and H4 protons have with the resonances in the 8.0-8.5 ppm range, which originate from multiple metabolites, including adenosine-containing compounds and NAA. The measurement of T_1_ and T_2_ relaxation times is necessary for absolute quantification of NAD^+^. In a subset of subjects, we found the NAD^+^ concentration to be 0.300 mM, determined from the H2 proton. This is in line with previous reports that measured NAD^+^ using 31P MRS (13). The studies to date that have quantified NAD^+^ concentration in human brain have used T_1_ and T_2_ correction factors extrapolated from the longer T_2_ measurement found at 11.7 T (12,15).

Because it is not feasible to measure metabolite T_1_ and T_2_ times in a subject-specific manner due to the lengthy nature of these experiments, it is typical to choose an estimate T_1_ and T_2_ (such the average T_1_ and T_2_ for NAD^+^ found in this study) to use as correction factors when calculating metabolite absolute concentrations. Doing so means that the uncertainty of the parameter estimates, described by the standard deviation of the measurements, propagates through to the final concentration estimate (30). This study thus additionally investigated the propagation of uncertainty for the T_1_ and T_2_ measurements for the H2 NAD^+^ proton. We found that the uncertainty of the T_2_ measurement has a much larger contribution to the final concentration uncertainty in comparison to the uncertainty of the T_1_ measurement by two orders of magnitude for the given protocol, as shown by the partial derivatives of concentration with respect to each relaxation time. The overall relative uncertainty given by the analytic approach was 8.5% while the uncertainty determined by the Monte Carlo simulation was 11.4%. This discrepancy may be explained in part by the fact that the analytic approach relies upon a first-order Taylor approximation and did not consider parameter dependencies, and a Monte Carlo approach could be more useful in exploring uncertainty propagation when the output function in nonlinear and results in a non-Gaussian distribution (32). In one previous study the intra-subject coefficient of variation of measured NAD^+^ concentration measured on the same day was 6% (12); in another study, the intra-subject coefficient of variation of NAD^+^ measured on subsequent days was 10% (33). Therefore, we found that the uncertainty in NAD^+^ quantification introduced by the inter-subject variation in T_1_ and T_2_ is within the approximate range of the intra-subject variability in NAD^+^ concentration measured using ^1^H MRS.

There are a number of limitations present in the study. Firstly, saturation recovery inherently has a smaller dynamic range than inversion recovery, which could reduce our ability to accurately measure T_1_. Additionally, our T_1_ fits all had R^2^ greater than 0.98 for both the H2 and H6 NAD^+^ peaks, indicating that the saturation recovery experiment with our chosen delay times was sufficient for measuring T_1_. Another limitation is the large volume. While this increased the detectability of NAD^+^, we were not able to determine regional differences in T_1_ and T_2_ within the brain which could be of interest in some pathologies. Furthermore, the parameter uncertainty propagation performed in this study constitutes just a small fraction of contributing sources of error in metabolite quantification. Additional sources of uncertainty may include, but are not limited to, accuracy of tissue segmentation and decreased reliability of spectral fitting in lower SNR scans (e.g. longer TE or shorter TS). Finally, our subjects were within the limited age range of 25-30 years old, and the measured T_1_ and T_2_ relaxation times may not be robust estimations for older or younger populations.

In conclusion, we found NAD^+^ T_1_ and T_2_ relaxation times of human brain measured in vivo at 7 T to be shorter than those previously reported in rat brain at 11.7 T and performed uncertainty propagation analysis for these parameters. These relaxation times can be used as correction factors for the absolute quantification of NAD^+^, as well as for protocol design in order to choose optimal TE and TR.

## Acknowledgements

Research reported in this publication was supported by the National Institute of Biomedical Imaging and Bioengineering of the National Institutes of Health under award Number P41EB029460, and by the National Heart, Lung, and Blood Institute of the National Institutes of Health under award Numbers R01HL137984, R01HL169378 and F31HL158217.

